# Epigenetic regulator BMI1 promotes fusion-positive rhabdomyosarcoma proliferation and constitutes a novel therapeutic target

**DOI:** 10.1101/2020.04.18.048355

**Authors:** Cara E. Shields, Sindhu Potlapalli, Selma M. Cuya-Smith, Sarah K. Chappell, Dongdong Chen, Daniel Martinez, Jennifer Pogoriler, Komal S. Rathi, Shiv A. Patel, John M. Maris, Robert W. Schnepp

## Abstract

Rhabdomyosarcoma (RMS) is an aggressive pediatric soft tissue sarcoma that continues to present significant challenges to pediatric oncology. There are two major subtypes of pediatric rhabdomyosarcoma, alveolar and embryonal. Alveolar rhabdomyosarcomas are characterized by the presence of a PAX-FOXO1 fusion protein and termed fusion-positive (FP-RMS); embryonal rhabdomyosarcomas (ERMS) lack these fusions and are termed fusion-negative (FN-RMS).

Fusion-positive rhabdomyosarcoma (FP-RMS) harbors PAX-FOXO1 fusion proteins and has a worse overall outcome compared to FN-RMS, underscoring the critical need to identify novel targets for this disease. While fusion proteins remain challenging therapeutic targets, recent studies have begun to reveal the key intersection of PAX-FOXO1 fusion proteins with the malignant epigenome, suggesting that epigenetic proteins may serve as novel targets in FP-RMS. Here, we investigate the contribution of the epigenetic regulator BMI1 to FP-RMS.

We examined RNA-seq tumor datasets and determined that *BMI1* is robustly expressed in FP-RMS tumors, patient derived xenografts (PDXs), and cell lines. We depleted BMI1 using RNA interference and find that this leads to a marked decrease in cell growth. Next, we used two BMI1 inhibitors, PTC-209 and PTC-028, and showed that BMI1 inhibition decreases cell cycle progression and increases apoptosis in FP-RMS cell lines. In the *in vivo* setting, targeting BMI1 leads to decreased tumor growth. Mechanistically, we observe that BMI1 inhibition activates the tumor suppressive Hippo pathway. Collectively, these results identify BMI1 as a novel therapeutic vulnerability in FP-RMS and provide a foundation for further investigation of BMI1 in both FP-RMS and additional sarcoma histotypes.

## INTRODUCTION

Rhabdomyosarcoma (RMS) is a tumor of developing skeletal myoblast-like cells that primarily afflicts children.^1^ There are two main subtypes of RMS, fusion positive (FP-RMS) and fusion negative (FN-RMS), which are classified by the presence or absence of the PAX-FOXO1 fusion protein.^1,2^ FP-RMS typically encompasses alveolar rhabdomyosarcoma (ARMS), with FN-RMS emerging as the preferred term for embryonal rhabdomyosarcoma (ERMS).^1^ These subtypes are based upon histological observations, but as we move more toward defining cancers molecularly, utilizing fusion status is more useful and accurate.^1^ FP-RMS has a worse outcome compared to FN-RMS, with an overall survival rate of below 30%, and an even more dire prognosis for patients with metastatic disease.^3^ Currently, the standard of care is multimodal and intensive, consisting of multiagent chemotherapy, radiation, and surgery.^4,5^ Given the substantial morbidity and mortality of FP-RMS, there is a need for novel, translatable treatment options.

While the PAX-FOXO1 fusion proteins are pathognomonic for this disease and contribute significantly to its aggression, they remain challenging drug targets.^1,6-8^ To date, efforts to inhibit PAX-FOXO1 directly have not yielded fruitful clinical results.^6^ Moreover, a recent study has suggested that PAX3-FOXO1 is necessary for the initiation/maintenance of FP-RMS but may not be needed in recurrence, suggesting that the targeting of diverse oncogenic networks may be necessary to optimize the treatment of this cancer.^7,9^ The interaction of PAX-FOXO1 fusions with the epigenome has become increasingly appreciated.^9-11^ PAX3-FOXO1 has been found to require BRD4 at superenhancers, suggesting a novel epigenetic vulnerability in FP-RMS.^10^ Further, the fusion protein requires CHD4, which is needed for chromatin remodeling, to activate a subset of its target genes.^12^ Histone deacetylases have also been investigated and found to control SMARCA4, which subsequently regulates *PAX3-FOXO1* mRNA stabilization.^13^ Clearly, these studies provide evidence for a significant relationship between the epigenome and the tumorigenicity of FP-RMS, and suggest the possibility that additional druggable epigenetic regulators may exist.

Inspired by these studies, we engaged in a search for additional druggable epigenetic complexes involved in FP-RMS. The Polycomb group proteins are epigenetic complexes traditionally associated with gene repression by chromatin compaction. They consist of Polycomb Repressive Complex 1 and 2 (PRC1/2) which control monoubiquitination of H2AK119 and trimethylation of H3K27, respectively.^14^ Dysregulation of PRC1/2 protein members are correlated with tumor initiation and progression in many adult cancers but remains relatively understudied in pediatric cancers.^15^ PRC2 members such as EZH2 have been analyzed and found to promote survival in the context of FP-RMS.^16^ Thus, in turn, we hypothesized that a member of PRC1, B lymphoma Mo-MLV insertion region 1 (BMI1) would be a viable epigenetic target in FP-RMS. BMI1 has no enzymatic activity itself but is a required component of PRC1 and is a known oncogene in numerous adult cancers including hematological malignancies, breast cancer, ovarian cancer, and more.^17-20^ BMI1 has also been studied in a few pediatric cancers, including glioblastoma and Ewing sarcoma, but remains unstudied in RMS.^21-23^ Additionally, BMI1 has been found to promote self-renewal in skeletal muscle and was also one of the components, along with TERT and PAX3-FOXO1, used to transform normal human myoblasts into a cell culture model of FP-RMS.^24,25^ In these studies, we identify BMI1 as a novel therapeutic liability in FP-RMS.

## MATERIALS & METHODS

### *In silico* data

The GTEx RNA sequencing data from 31 normal adult tissues comprising 7,863 samples was downloaded from S3 buckets (Amazon; s3://cgl-rnaseq-recompute-fixed/target/ and s3://cgl-rnaseq-recompute-fixed/gtex/) on 8/5/2016 from prior processed data as described from the UCSC Computational Genomics laboratory (Vivian et al., 2016). RNA-sequencing data of 15 RMS patient-derived xenograft (PDX) models from the Pediatric Preclinical Testing Consortium (PPTC) was processed using the STAR alignment tool and subsequently normalized using the RSEM package based upon the hg38 reference genome and the GENCODE v23 gene annotation. Gene expression values were quantified in terms of Fragments Per Kilobase per Million mapped reads (FPKM).

### Cell culture

Rhabdomyosarcoma cell lines (Rh30 and Rh41) were obtained from the Children’s Hospital of Philadelphia (Courtesy of Dr. Margaret Chou) as well as from the Children’s Oncology Group (Rh28 and CW9019). The Emory Genomics Core authenticated cell lines for use and *Mycoplasma* testing was performed every 3 - 6 months using the *Mycoplasma* test kit (PromoCell, PK-CA91-1024). Cells were cultured in a humid incubator at 37°C with 5% CO2. Rh30 and CW9019 were passaged regularly in DMEM (Corning) and Rh28 and Rh41 were passaged in RPMI 1640 (Corning). Media was supplemented with 10% FBS (Corning) and 1% L-glutamine (Gemini). No antibiotics or antimycotics were added to the media.

### Plasmids and lentiviral preparation

BMI1 shRNA plasmids were purchased from Sigma (pLKO.1). The catalog numbers are shBMI1-2: TRCN0000020156 and shBMI1-4: TRCN0000218780. Generation of infectious lentiviral particles and subsequent cell transduction was performed as previously described.^26^ FuGENE 6 (Promega) was used to transfect select plasmids, with pMD2.G (VSV-G plasmid) and psPAX2 (packaging plasmid), into HEK293T cells. Viral supernatant was collected 2-3 days after transfection and filtered with a 0.45 µm nitrocellulose membrane. Following this, cells were transduced with viruses. One million cells were seeded into 10 cm plates and transduced, along with 8 µg/mL polybrene (Sigma). Fresh media was added 6 hr post virus addition. The next day, the media was replaced completely with fresh media. Two days later, puromycin was added to select for transduced cells.

### siRNA transfection

Cells were plated at 200,000 cells per well in a 6 well plate. The following day, cells were transfected using DharmaFECT 1 (Horizon Discovery) and 25 nM of an siRNA ON-TARGET Plus SMARTpool (Horizon Discovery) or ON-TARGET Plus Non-targeting Control Pool (Horizon Discovery). Cells were harvested for analysis 72 hr post-transfection.

### Real-Time PCR and Western blots

RNA was isolated from cells using the RNeasy Mini Kit (QIAGEN) and Real-Time PCR (RT-PCR) analysis performed as previously described.^26^ For western blots, cell samples were lysed in RIPA (Boston Bioproducts) containing cOmplete protease inhibitor cocktail (Roche) and PMSF (Cell Signaling Technology) then sonicated. Protein concentrations were determined using the Bradford assay (Bio-Rad) and samples (20 μg protein) run on SDS PAGE Bis-Tris 4-12% gels (Life Technologies). The gels were transferred to nitrocellulose membranes and membranes blocked in 5% Blotting-Grade Blocker (Bio-Rad) in Tris-Buffered Saline with 1% Tween-20 (Cell Signaling Technology). The blots were incubated with primary antibodies in 5% BSA (Jackson Laboratory) overnight at 4°C. The secondary antibodies used were IRDye 800CW/680RD anti-Rabbit or anti-Mouse (Li-COR Biosciences) at 1:50,000 and 1:5,000, respectively. Whole blots were scanned using the Li-COR Odyssey. The primary antibodies and dilutions are listed in Supplementary Table 1. Any quantifications are presented as relative adjusted densities and were performed in ImageJ.

### Cell growth assays

CellTiter-Glo (Promega) and Caspase-Glo (Promega) were used to assess viability of both shRNA/siRNA manipulated and drug treated cells. On day 0, 2,000 cells/well were plated in a 96 well plate and on day 1 treated with control or drug. To calculate IC50s, cells were treated with a 7-log dose range of inhibitor (10^−11^M - 10^−5^M). Cells proliferated for an additional 96 hr before performing CellTiter-Glo or Caspase-Glo per the manufacturer’s instructions. IC50s were calculated by log transforming concentrations, fitting to a three-parameter logistic nonlinear regression curve and finding the half-maximal concentration.^27^

For crystal violet colony formation assays, we plated 2,000 cells/well in duplicate in 6 well plates. We treated cells with drugged media and allowed cells to proliferate for 10 days prior to washing/fixing with 3.7% formaldehyde then staining with 0.0025% crystal violet. Plates dried overnight and were imaged with a Nikon D3400.

### Flow cytometry

On day 0, cells were seeded at 1 million cells /10 cm plate and PTC-028 added on day 1. Cells were harvested after 48 hr. Staining was performed using Annexin V-FITC/PI (BD Biosciences) or BrdU-APC/7-AAD (BD Biosciences) kits following manufacturer’s instructions. For Annexin V/PI staining, cell media containing dead cells in suspension was also collected. Samples were run within 1 hr on a Cytoflex 96 well plate loader, with 50,000 - 100,000 events collected per sample. Compensation, gating and analyses were performed in FlowJo.

### *In vivo* xenograft model

Heterozygous nude mice (Crl:NU(NCr)-*Foxn1*^*nu*^*/*^*+*^) between 5 - 6 weeks old (Charles River) were housed in sterile cages at the Health Sciences Research Building Animal Facility at Emory University. Mice acclimated to their new environment for 1 week after being received and were maintained in 12 hr day/night cycles. All experimental procedures were Emory IACUC approved. 2 million Rh30 cells were mixed 1:1 with Matrigel (Corning) and subcutaneously injected into the right flank of each mouse. As previously described, treatments began when tumors were equal to or greater than 100 mm^3^.^28,29^ The mice were tagged and randomly separated into 2 groups: vehicle (n = 10) and PTC-028 (n = 10). Mice received vehicle (0.5% HPMC, 1% Tween-80) or 15 mg/kg PTC-028 twice weekly by oral gavage.^28,29^ Weights and tumor sizes were measured three times weekly. Tumor volumes were calculated by using an ellipsoid volume formula: π / 6 x L x W x H.^30^ In accordance with the IACUC protocol, mice were sacrificed when tumors reached a volume greater than or equal to 1500 mm^3^. Collected tumors were removed post-mortem and snap-frozen in liquid nitrogen for immunoblotting or formalin fixed and paraffin embedded for immunohistochemistry.

### Immunohistochemistry

A tumor array of pediatric sarcomas (duplicate punches) was constructed at The Children’s Hospital of Philadelphia. An additional normal pediatric tissue array consisted of duplicate punches of 41 normal pediatric tissues/organs procured from the Children’s Hospital of Philadelphia from 2005 – 2012. BMI1 antibody (Cell Signaling Technology) was used to stain formalin fixed paraffin embedded tissue slides. Staining was performed on a Bond Rx automated staining system (Leica Biosystems). The Bond Refine polymer staining kit (Leica Biosystems) was used. The standard protocol was followed apart from the primary antibody incubation which was extended to 1 hr at room temperature and the post primary step was excluded.^27^ BMI1 antibody was used at a 1:200 dilution and antigen retrieval was performed with E1 (Leica Biosystems) retrieval solution for 20 min. Slides were rinsed, dehydrated through a series of ascending concentrations of ethanol and xylene, then coverslipped. Stained slides were then digitally scanned at 20x magnification on an Aperio CS-O slide scanner (Leica Biosystems).

### Statistical analyses

Data analyses were performed in GraphPad Prism 8. Statistical significance was determined using an unpaired student two tailed t-test for two groups. Groups of three or more were analyzed using an ANOVA. All assays were performed in duplicate unless otherwise stated and presented using mean and standard deviation. Survival curves were generated in Prism 8 using the Kaplan-Meier method.^31^

## RESULTS

### BMI1 is highly expressed in rhabdomyosarcoma

To investigate BMI1 as a potential therapeutic vulnerability in FP-RMS, we sought to define its expression pattern in sarcomas, broadly considered. We first examined Oncomine and determined the expression of *BMI1* in both adult and pediatric sarcomas.^32^ We noted that *BMI1* is robustly expressed in pediatric sarcomas, such as Ewing sarcoma and osteosarcoma, as well as in adult subtypes, including leiomyosarcoma and chondrosarcoma (Supp. Fig. S1A - B).^22,32,33^

Next, we focused on RMS. We began by interrogating available datasets and first looked at human exon array data from both FP-RMS and FN-RMS patient tumor samples.^34^ We observed that *BMI1* expression levels were robustly expressed across both subtypes (Fig. 1A). To focus on FP-RMS specifically, we compared *BMI1* levels from RNA-seq FP-RMS patient-derived xenograft (PDX) data from the Pediatric Preclinical Testing Consortium (PPTC) to normal tissues (GTEx).^35^ We found that *BMI1* mRNA levels are highly expressed in FP-RMS compared to normal tissues (Fig. 1B). Furthermore, we probed the OncoGenomics database and found *BMI1* to be highly expressed in both FP-RMS and FN-RMS (Supp. Fig. S1C).^36^ We performed a tumor microarray with FP-RMS patient samples and confirmed that BMI1 is robustly expressed at the level of protein (Fig. 1C).

**Figure 1.**
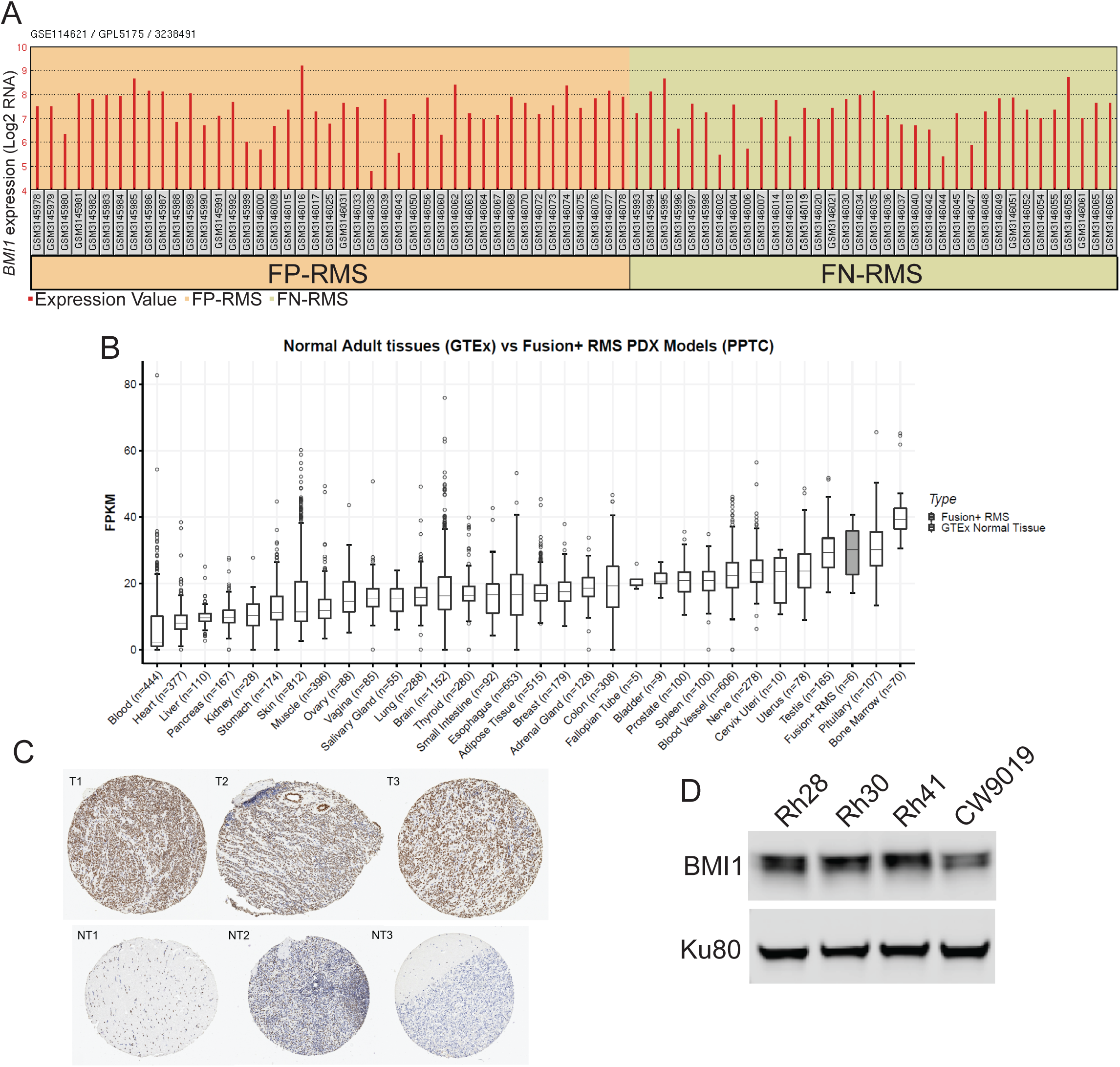
BMI1 is highly expressed in rhabdomyosarcoma. (A) Barplot of *BMI1* gene expression (Log2 RNA signal intensity) from human exome array data across FP-RMS and FN-RMS patient tumor samples (GSE114621).^34^ (B) Boxplot of *BMI1* gene expression values from RNA-sequencing data across GTEx normal tissues (n = 31) and FP-RMS PDX models (n = 6). Y-axis represents FPKM values. (C) Tumor microarray with three patient FP-RMS tumors (T1, T2, T3), compared to normal pediatric tissue (NT1 = pediatric skeletal muscle, NT2 = pediatric spleen, NT3 = pediatric cerebellum). BMI1 is brown (DAB). The nuclear counterstain for BMI1-negative cells is purple (hematoxylin). (D) Western blot of FP-RMS cell lines Rh28, Rh30, Rh41 and CW9019 showing BMI1 protein expression with a Ku80 loading control.

Finally, we surveyed the expression of BMI1 across the FP-RMS cell lines Rh28, Rh30, Rh41 and CW9019 and find that BMI1 is robustly expressed across all models (Fig. 1D). Notably, Rh28, Rh30, and Rh41 have the PAX3-FOXO1 fusion, while CW9019 harbors the PAX7-FOXO1 fusion.

### Genetic knockdown of BMI1 leads to reduced cellular proliferation in FP-RMS cells

Our analyses demonstrate that BMI1 is highly expressed in both fusion-positive and negative rhabdomyosarcoma. Given the clinical aggression of FP-RMS, in subsequent investigations, we focused exclusively on this subtype. First, we depleted BMI1 using two independent shRNAs directed against BMI1 and confirmed effective knockdown of BMI1 (Fig. 2A - B). In two FP-RMS cell line models (Rh28 and Rh30), we observed that BMI1 knockdown significantly reduces cell proliferation (Fig. 2A - B). Knockdown of BMI1 was confirmed by Western blot (Fig. 2A - B). To further validate these findings, we utilized pooled siRNAs (comprised of 4 independent siRNAs directed against BMI1) to transiently deplete BMI1 and again demonstrated significantly decreased proliferation (Fig. 2C - D). Knockdown of *BMI1* was confirmed by RT-PCR (Fig. 2C - D) These results suggest that BMI1 promotes cell proliferation in FP-RMS.

**Figure 2.**
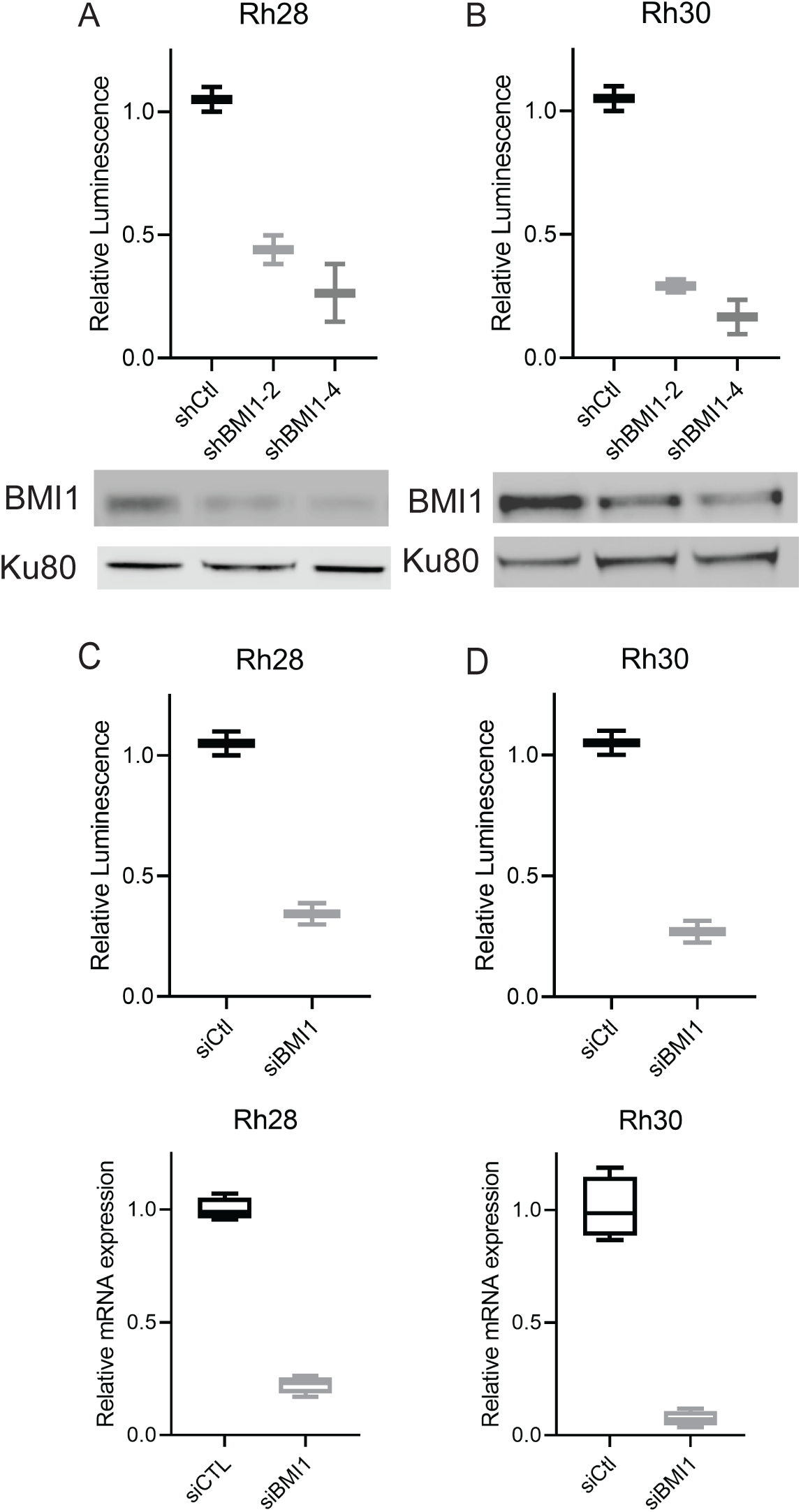
Genetic knockdown of BMI1 leads to reduced cellular proliferation in FP-RMS cells. (A-B) Rh28 (A) and Rh30 (B) cell lines were infected with control lentiviruses or lentiviruses expressing two independent shRNAs directed against BMI1. Cell proliferation in control and BMI1-depleted cell lines as assessed by Cell-TiterGlo. Western blotting of BMI1 and Ku80 in corresponding cell lines. (C-D) Rh28 (C) and Rh30 (D) cells were transfected with control siRNAs or pooled siRNAs directed against BMI1. Cell proliferation assessed by Cell-TiterGlo, with corresponding siCtl and siBMI1 RT-PCR data depicted below. Standard deviation bars shown. Results are representative of at least three independent experiments.

### Pharmacologic inhibition of BMI1 decreases cell proliferation *in vitro*

We next assessed the effects of pharmacologic inhibition of BMI1 on FP-RMS. To do so, we initially employed PTC-209, an inhibitor that reduces BMI1 protein levels and lowers PRC1 activity in cancer cells, with minimal effects in non-cancerous cell line models.^37^ In several aggressive cancer models, such as colorectal cancer and biliary tract cancer, PTC-209 has been found to impair cell growth through promoting cell cycle arrest and causing cell death.^37,38^ Guided by previous studies, we treated 4 FP-RMS cell lines with PTC-209 across a 7-log dose range (10^−11^ M - 10^−5^ M). Treatment with PTC-209 significantly decreases cell proliferation (Fig. 3A - D) in all 4 cell lines, with IC50s ranging from 483 nM to 872 nM (Fig. 3K). Protein levels of BMI1 were also reduced with PTC-209 treatment (Supp. Fig S2A).

**Figure 3.**
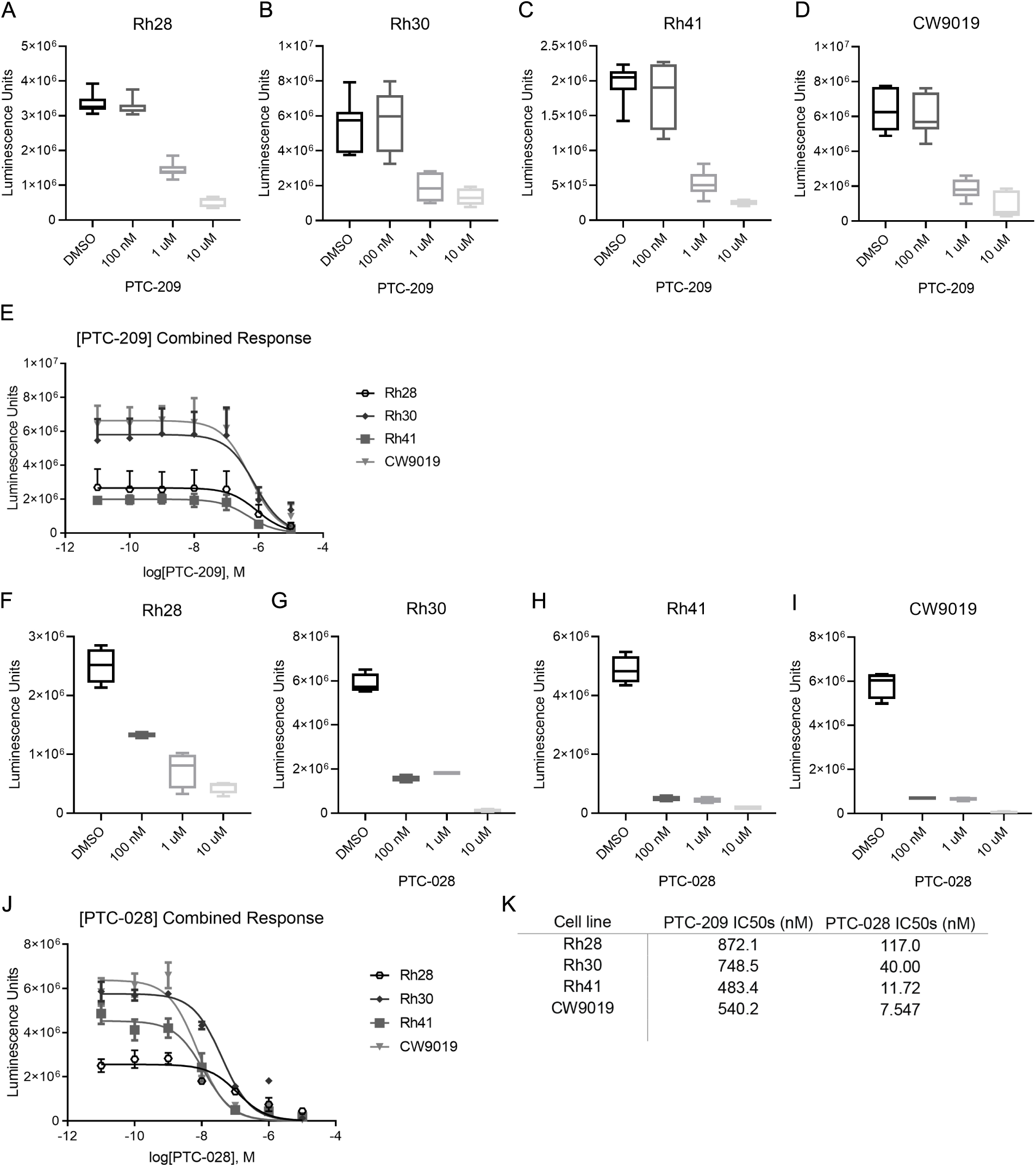
Pharmacologic inhibition of BMI1 decreases cell proliferation *in vitro*. (A-D) Cell lines Rh28 (A), Rh30 (B), Rh41 (C) and CW9019 (D) were treated with a 7-log dose range of PTC-209. Graphs display cell viability measured with CellTiter-Glo with varying concentrations of PTC-209. E. Dose response curve of PTC-209 ranging from 10^−11^ M – 10^−5^ M. (F-I) Cell lines Rh28 (F), Rh30 (G), Rh41 (H) and CW9019 (I) were treated with a 7-log dose range of PTC-028. Graphs display cell viability measured with CellTiter-Glo at varying concentrations of PTC-028. (J) Dose response curve of PTC-028 ranging from 10^−11^ M – 10^−5^ M. (K) Table summarizing IC50 values of PTC-209 and PTC-028. Standard deviation bars depicted. Results are representative of at least three independent experiments.

Next, we assessed the impact of a second generation BMI1 inhibitor, PTC-028, on FP-RMS proliferation. PTC-028 inhibits BMI1 by a different method than PTC-209, resulting in hyperphosphorylation of BMI1 and disrupting its function.^28^ It is also orally bioavailable, allowing for preliminary investigation of BMI1 disruption in the *in vivo* setting; for these reasons, in subsequent studies we employed PTC-028. Treatment with PTC-028 similarly decreases cell proliferation (Fig. 3F - J) in all 4 cell lines, yielding decreased BMI1 protein levels (Supp. Fig S2A). As expected, IC50s were lower for PTC-028 than for PTC-209, consistent with the greater potency of PTC-028 (Fig. 3K). Additionally, brightfield microscopy and colony formation assays showed that viability is significantly diminished with 50 nM and 100 nM doses of PTC-028 in Rh30 and CW9019 (Supp. Fig S2B - C). Thus, our data indicate that two BMI1 inhibitors greatly decrease proliferation in FP-RMS cell line models, mimicking the effects we observed with genetic disruption of BMI1.

### Targeting BMI1 decreases cell cycle progression and increases apoptosis in FP-RMS

We next aimed to define the mechanisms by which BMI1 promotes cell proliferation. Previous investigations have demonstrated that BMI1 influences cell cycle progression in part through repression of the *CDKN2A* (*p16-INK4a*) locus^39^, although this regulation is not observed in all contexts. BMI1 also possesses functions independent of *CDKN2A* repression, including the regulation of genes involved in differentiation and cell contact inhibition in Ewing sarcoma and androgen receptor expression in prostate cancer^22,40^.

To investigate the influence of BMI1 on cell cycle progression, we treated Rh30 with PTC-028 at doses below and near the IC50 of Rh30 and then performed BrdU/7-AAD staining. We observed an increase in the sub-G1 population and a decrease in the percentage of cells in S phase when the cells were treated with 50 nM of PTC-028 for 24 hr (Fig. 4A-4B). Given the increase in the sub-G1 population, we speculated that BMI1 additionally increases apoptosis *in vitro*. Therefore, we performed Annexin V/PI staining and observed a dose-dependent increase in the percentage of apoptotic cells (Fig. 4C - 4D). To further verify the apoptosis phenotype, we probed for cleaved PARP and noted an increase in PARP cleavage with PTC-028 addition (Fig. 4E). Additionally, to complement these data, we performed Caspase-Glo analyses of shBMI1/siBMI1 Rh28 and Rh30 cell lines and discovered an increase in caspase 3/7 activity (Supp. Fig S3A - B). We delved down further and analyzed apoptosis in siBMI1 transfected Rh28 and Rh30 cells by Annexin V/PI staining and again noted an increase in the apoptotic fractions (Supp. Fig S3C). Together, these data confirm that pharmacologically targeting BMI1 impairs progression to S phase and results in apoptosis.

**Figure 4.**
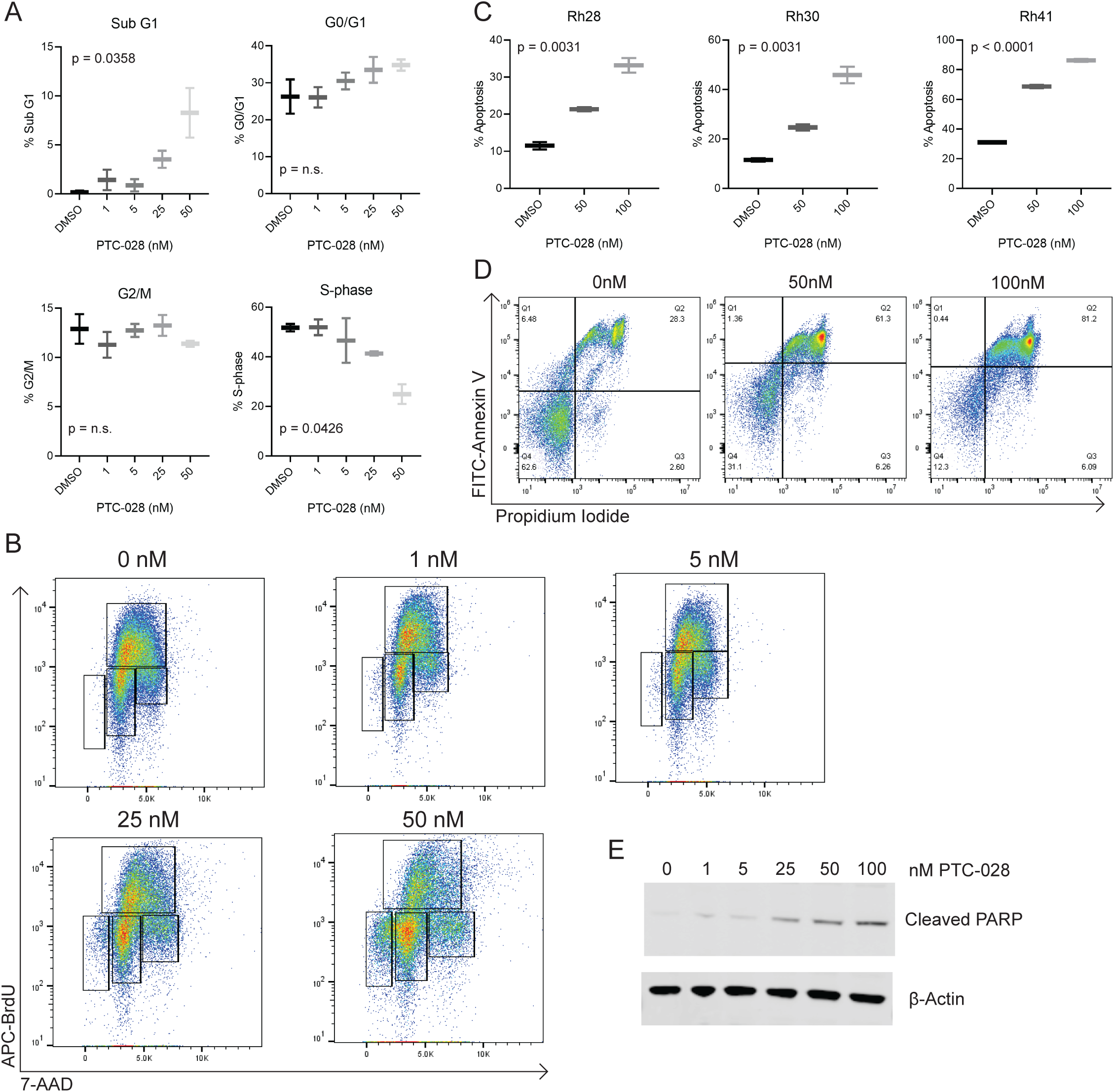
Targeting BMI1 decreases cell cycle progression and increases apoptosis in FP-RMS. (A) Graphs depict cell cycle distribution in the Rh30 cell line treated with PTC-028 (0 - 50 nM). (B) Representative cell cycle distribution from Rh30. BrdU is depicted on the y-axis with 7-AAD on the x-axis. (C) Flow cytometry analysis of Annexin V/PI staining in Rh28, Rh30 and Rh41, with PTC-028 treatment ranging from 0 - 100 nM. (D) Representative example of flow cytometry data illustrating apoptosis with Annexin V (y-axis) and propidium iodide (x-axis). (E) Rh30 was treated with PTC-028 for 72 hr, with Western blot depicting cleaved PARP and actin. Standard deviation bars depicted. Results are representative of at least three independent experiments.

### Single agent PTC-028 treatment causes tumor growth delay *in vivo*

To provide the initial foundation for targeting BMI1 in FP-RMS, we employed PTC-028, which is orally bioavailable.^28,29^ Nude mice bearing Rh30 xenografts were treated with vehicle or PTC-028 (15 mg/kg by oral gavage) daily, a dosing scheme guided by previous studies^28,29^. As shown in Fig. 5A, treatment with PTC-028 delays tumor growth in comparison to vehicle (Fig. 5A, p = 0.0005). The treatment was well-tolerated, with no significant change in weights (Fig. 5B) and no signs of pain or distress in the mice observed. The vehicle group died by day 25, while the PTC-028 treated group survived until day 41 (Fig. 5C, p = 0.0002). The tumors were harvested and analyzed for BMI1 protein levels. By Western blot, we noted that tumors in PTC-028 treated mice had an approximately 30% reduction in BMI1 levels in comparison to control. (Fig. 5D). Interestingly, however, in contrast to the *in vitro* setting, we noted no increase in cleaved PARP (Fig. 5E). Collectively, these results suggest that single-agent treatment with PTC-028 delays, though does not abrogate, the growth of a FP-RMS xenograft.

**Figure 5.**
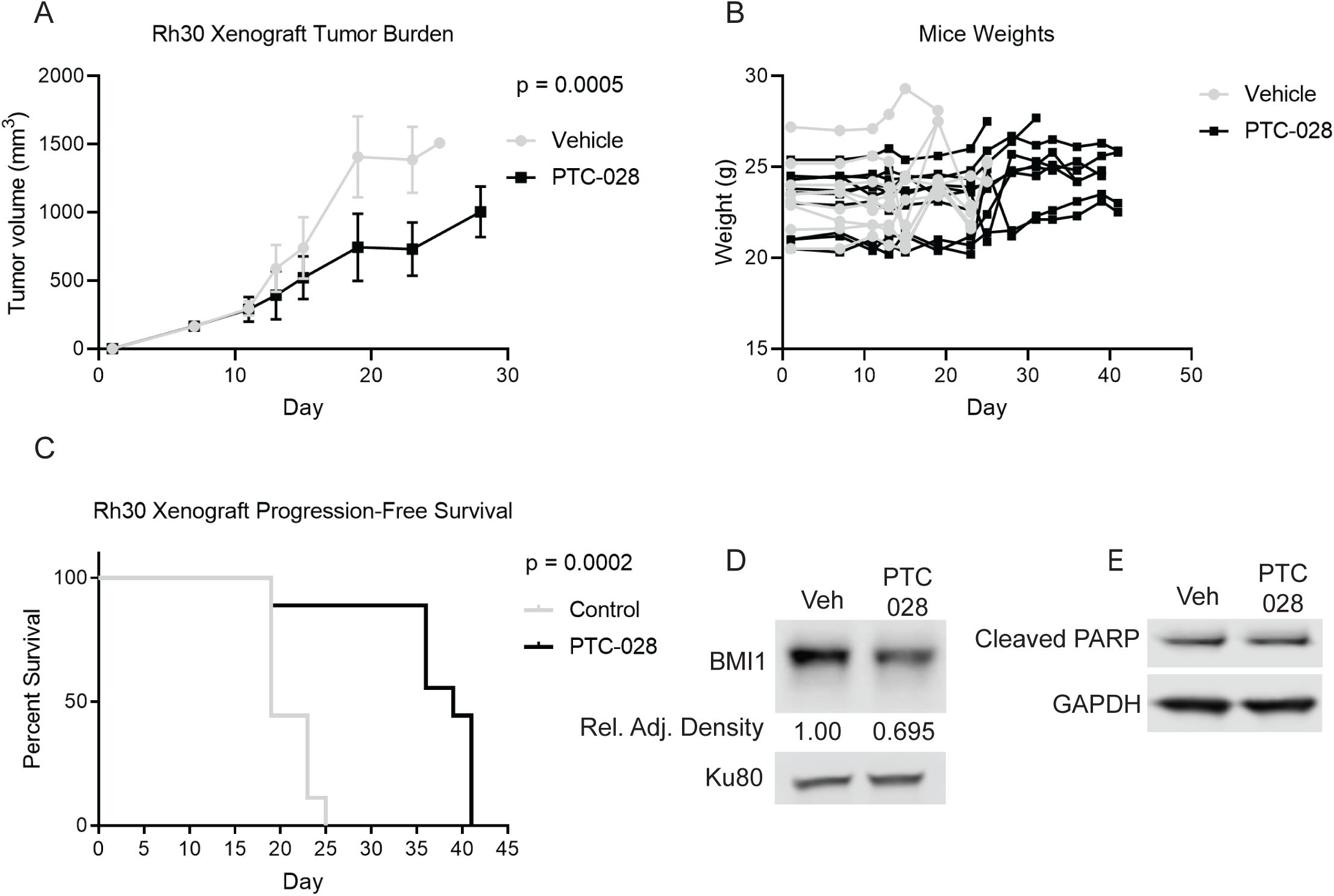
Single agent PTC-028 treatment causes tumor growth delay *in vivo*. Rh30 xenografts were treated with vehicle or PTC-028 (15 mg/kg 2x/weekly). (A) Response of tumor volumes to vehicle and PTC-028. (B) Weight change from baseline on study arms. (C) Kaplan-Meier analyses for Rh30 xenografts. (D) Representative Western blot of BMI1 and Ku80 in control and PTC-028 treated tumors. (E) Western blot of cleaved PARP levels with GAPDH as a loading control. Standard deviation bars are included.

### BMI1 negatively influences Hippo signaling

Given our findings demonstrating the positive influence of BMI1 on cell cycle progression, we first asked whether BMI1 inhibits *CDKN2A* expression in FP-RMS.^41^ A canonical target of BMI1 is *CDKN2A*, and repression of *CDKN2A* controls cell cycle progression to S phase.^39,41^ We found that BMI1 inhibition by PTC-028 treatment leads to a slight upregulation in CDKN2A protein levels (Supp. Fig. S4A).

We next undertook a candidate-based approach to identify additional novel BMI1-influenced signaling networks in FP-RMS. We focused on Hippo signaling as BMI1 has been reported to interact with the Yes-Associated Protein (YAP) in Ewing sarcoma.^22^ In addition, PAX3-FOXO1 has been found to suppress the Hippo pathway in FP-RMS, and loss of Hippo signaling by MST1 knockout was shown to accelerate FP-RMS tumorigenesis.^42,43^

We began with determining the effects of BMI1 inhibition on canonical Hippo signaling. Normally, YAP/TAZ binds TEAD and YAP/TAZ/TEAD complexes influence genes implicated in cell cycle progression and growth (Fig. 6D).^44^ MST1 phosphorylates and activates LATS1/2, which in turn phosphorylates YAP/TAZ, leading to YAP/TAZ degradation and subsequent reduction in the amount of YAP/TAZ/TEAD complexes.^44^ Upon treatment with PTC-028, we observed that LATS1/2 phosphorylation increases, and YAP levels decrease (Fig. 6A - 6B), suggesting that the Hippo pathway is activated when BMI1 is inhibited. However, there is no increase in MST1 phosphorylation (Supp. Fig. S4B - C), suggesting a possible alternative mechanism for the increase in LATS1/2 phosphorylation. We depleted BMI1 using siRNAs and similarly observed an increase in LATS1/2 phosphorylation and a decrease in YAP protein expression (Fig. 6C). Overall, BMI1 inhibition appears to promote Hippo pathway activation through LATS1 phosphorylation.

**Figure 6.**
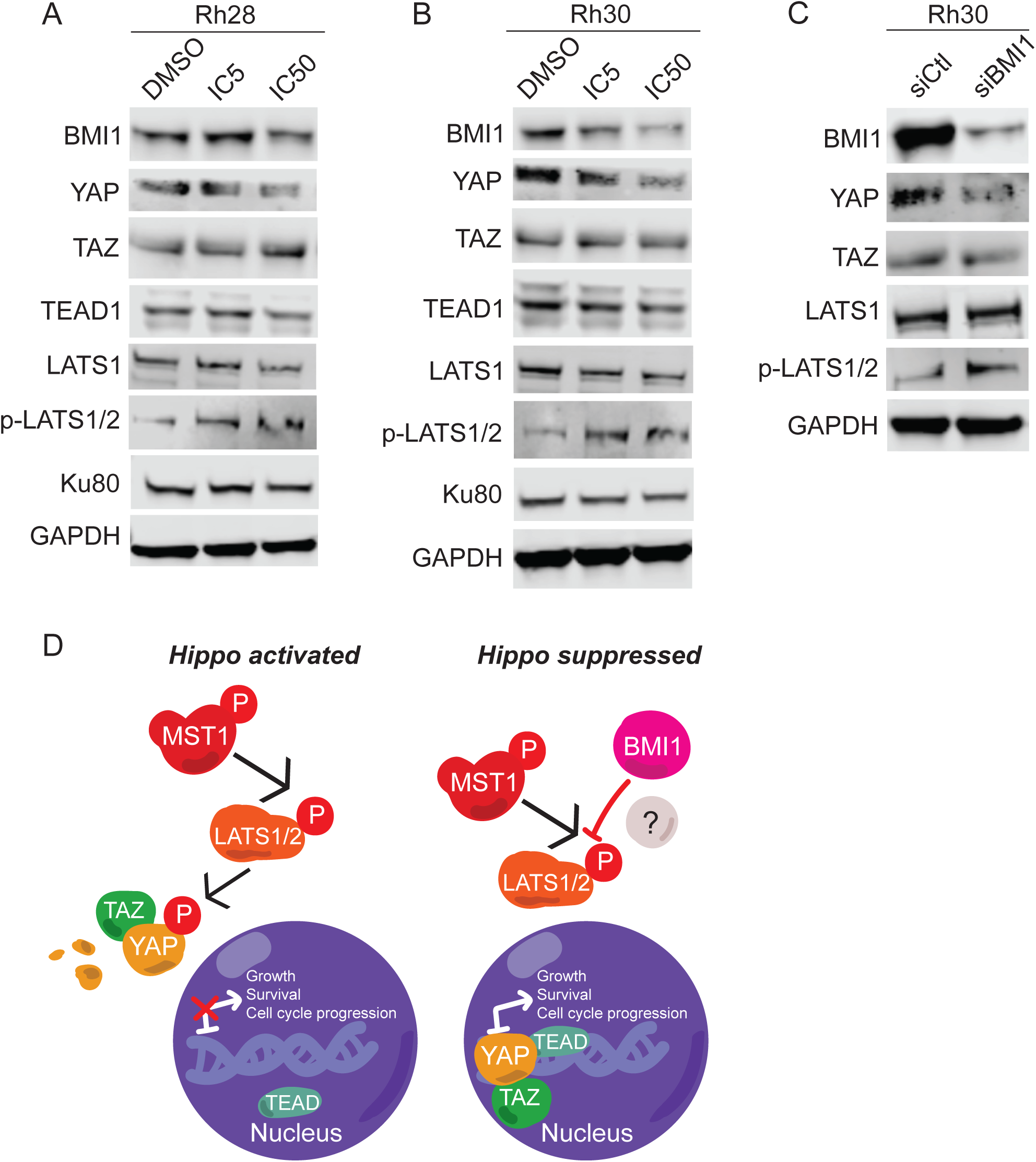
BMI1 negatively influences Hippo signaling. (A) Rh28 and (B) Rh30 cells were treated with PTC-028 at respective IC5 or IC50 concentrations for 72 hr, with DMSO as a control. Western blot of BMI1 and Hippo pathway members YAP, TAZ, TEAD1, LATS1, p-LATS1/2, and Ku80/GAPDH as loading controls. (C) Rh30 cells were transfected with an siRNA pool against BMI1 and Western blot analyses were performed after 72 hr. Western blot of BMI1 and Hippo pathway members YAP, TAZ, LATS1, p-LATS1/2, and GAPDH as loading controls. (D) Potential model of BMI1 involvement in the Hippo pathway. Results are representative of at least three independent experiments.

## DISCUSSION

Our understanding of, and hence optimal treatment for FP-RMS, remains inadequate. Motivated by a growing understanding that PAX3-FOXO1 fusion proteins interact with diverse epigenetic complexes, including BRD4^10,11^ and CHD4^12^, we hypothesized that BMI1 would contribute to FP-RMS aggression and that inhibiting this protein could potentially confer therapeutic benefit. Importantly, while studies suggest that BMI1 inhibition is a downstream effect of PTC-028^21^, our studies show that genetic depletion of BMI1 using multiple independent siRNAs/shRNAs diminishes proliferation (Fig. 2). Moreover, we find that pharmacologic disruption using PTC-209, which inhibits effective translation of *BMI1* mRNA^37^, decreases FP-RMS cellular viability significantly (Fig. 3). We provide evidence that BMI1 inhibition diminishes cell cycle progression and increases apoptosis (Fig. 4).

In the *in vivo* setting, we show that single agent treatment significantly decreases, though does not abrogate, FP-RMS growth (Fig. 5). Notably, while PTC-028 displays better *in vivo* characteristics than PTC-209, PTC-028 is still an early generation-inhibitor. PTC-596 is the clinical analog of PTC-028 that has recently entered into clinical trials for patients with advanced solid malignancies.^45^ A1016 is an additional BMI1 inhibitor related to PTC-596 and has shown similar positive results in glioblastoma.^21^ Future investigations will investigate the impact of these newer generation inhibitors on FP-RMS. Recently, investigators showed that the combination of PTC-596 and standard chemotherapy (gemcitabine and nab-paclitaxel) resulted in regressions in multiple aggressive pancreatic cancer models and, importantly, was well-tolerated.^46^ Based on such studies, we speculate that combining BMI1 inhibition with standard-of-care chemotherapeutic regimens in RMS may both be well-tolerated and result in greater inhibition of tumor growth, though further studies are needed to investigate this hypothesis.

While the current study delineates the impact of BMI1 on cell cycle progression and evasion of apoptosis, BMI1 has been implicated in multiple hallmarks of cancer, including DNA repair and self-renewal, among others.^39^ In melanoma, *BMI1* expression was shown to be correlated with an invasive signature and to promote multiple aspects of melanoma metastasis, including anoikis, invasion, migration, and chemoresistance.^47^ Might BMI1 contribute to metastatic dissemination in FP-RMS and could disruption of its function impede metastatic dissemination? Finally, while our studies focused on FP-RMS, we find that *BMI1* is broadly expressed in multiple pediatric and adult sarcomas (Fig. 1). It will be of interest to investigate the effects of BMI1 on the initiation, maintenance, and progression of various sarcomas.

In addition to proposing a role for BMI1 in FP-RMS aggression, our studies also reveal the influence of BMI1 on Hippo signaling and raise further mechanistic questions. For example, we find that inhibition of BMI1 results in increased levels of LATS1/2 phosphorylation at Thr1079/Thr1041, which is associated with LATS1/2 activation.^48^ However, inhibiting BMI1 does not appear to influence either the expression or phosphorylation of MST1, which lies upstream of LATS1 (Fig. 6). It is possible that BMI1 normally epigenetically represses an unidentified kinase of LATS1, or perhaps BMI1 engages with LATS1 through protein-protein interactions (Fig. 6D). Further investigation is necessary to define the mechanism of action by which BMI1 influences Hippo signaling. Interestingly, in undifferentiated pleomorphic sarcomas, there is evidence for the deregulation of the Hippo pathway and subsequent activation of YAP/TAZ.^49^ It is intriguing to posit a broad role for BMI1 involvement in the Hippo pathway across sarcomas and to speculate that BMI1 inhibition may provide a method of activating the Hippo pathway in these malignancies.

In conjunction with further dissection of BMI1-Hippo signaling, it will be important to define the full repertoire of genes influenced by BMI1 using both RNA-seq and ChIP-seq approaches, and to see how BMI1-influenced genes converge and diverge from other malignancies.^21,40,50^ Furthermore, it will be of substantial interest to determine if BMI1 acts through its canonical role as a member of the PRC1 complex, or by associating with other complexes to control gene expression in FP-RMS. Moreover, what effects does BMI1 inhibition have on global chromatin changes? Additional ChIP-seq experiments investigating the impact of BMI1 inhibition on histone repressive marks such as H2AK119Ub and H3K27me3, along with active marks like H3K27ac, will help clarify the molecular mechanisms by which BMI1 influences the malignant phenotype.

Our studies propose a novel role for BMI1 signaling in FP-RMS, connect BMI1 to Hippo signaling, and raise additional questions with regards BMI1 function and signaling. Finally, they provide an initial foundation for investigating the utility of BMI1 inhibition in FP-RMS and perhaps additional sarcoma subtypes.

## AUTHOR CONTRIBUTIONS

Conception and design: C.E. Shields, R.W. Schnepp

Development of methodology: C.E. Shields, R.W. Schnepp

Acquisition of data: C.E. Shields, S. Potlapalli, S.M. Cuya, S.K. Chappell, D. Chen, D. Martinez, J. Pogoriler, S. Patel, R.W. Schnepp

Analysis and interpretation of data (biostatistics, statistical analysis, interpretation of clinical data and genomic datasets): C.E. Shields, K.S. Rathi, R.W. Schnepp

Writing, review and/or revision of the manuscript: C.E. Shields, R.W. Schnepp

Administrative, technical, or material support: J.M. Maris, R.W. Schnepp

Study supervision: R.W. Schnepp

## ACKNOWLEDGEMENTS

This work was supported in part by NIH Grant K08-7K08CA194162-02 (R.W.S), NIH Grant R35 CA220500 (J.M.M.), the Sarcoma Foundation of America (R.W.S), CURE Childhood Cancer (R.W.S), Austen’s Army (R.W.S), the Aflac Cancer and Blood Disorders Center Trust (R.W.S), and the William Woods, M.D., Aflac Clinical Investigator Chair (R.W.S.).

Additionally, this study was supported in part by the Emory Flow Cytometry Core (EFCC), one of the Emory Integrated Core Facilities (EICF), and is subsidized by the Emory University School of Medicine. Additional support was provided by the National Center for Georgia Clinical & Translational Science Alliance of the National Institutes of Health under Award Number UL1TR002378. The content is solely the responsibility of the authors and does not necessarily represent the official views of the National Institutes of Health.

