## Supplementary Figures for "Epigenetic regulator BMI1 promotes fusion-positive rhabdomyosarcoma proliferation and constitutes a novel therapeutic target"

### Supplementary Figure S1

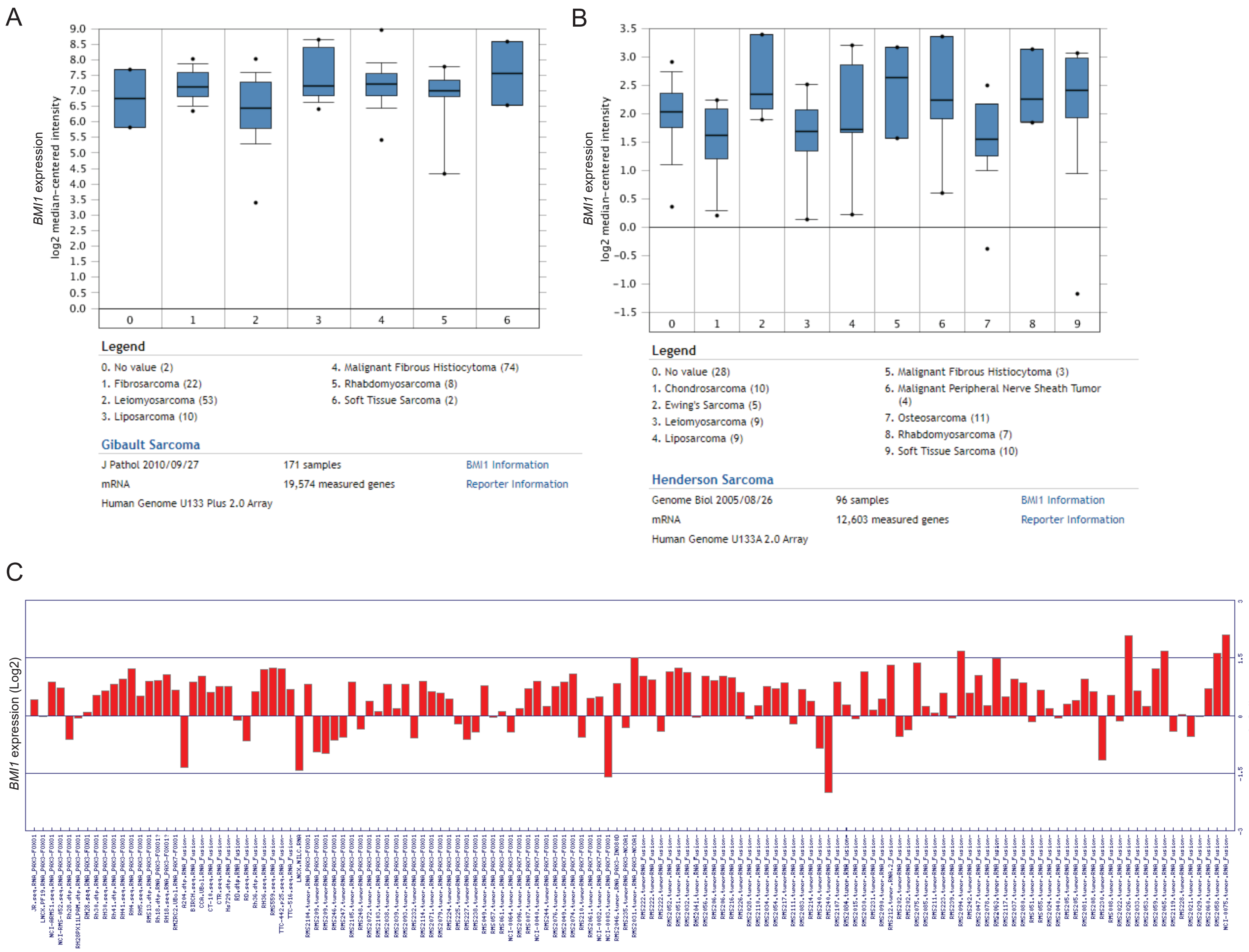

### Supplementary Figure S2

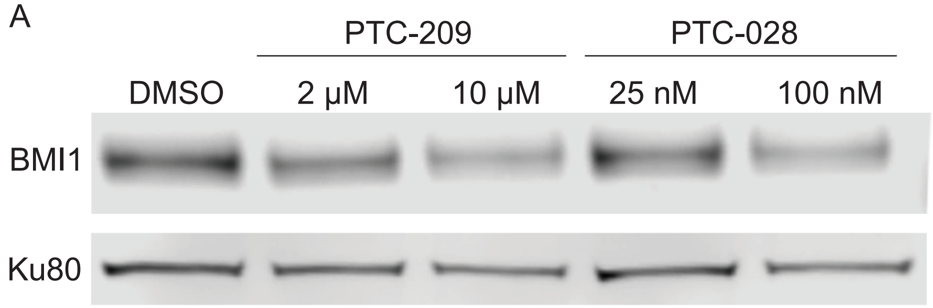


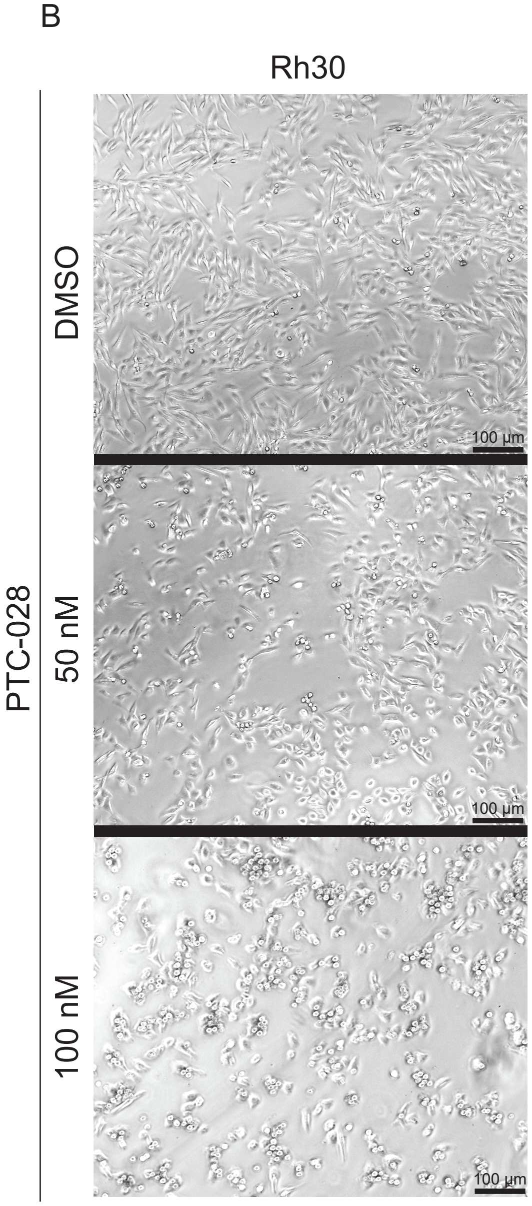


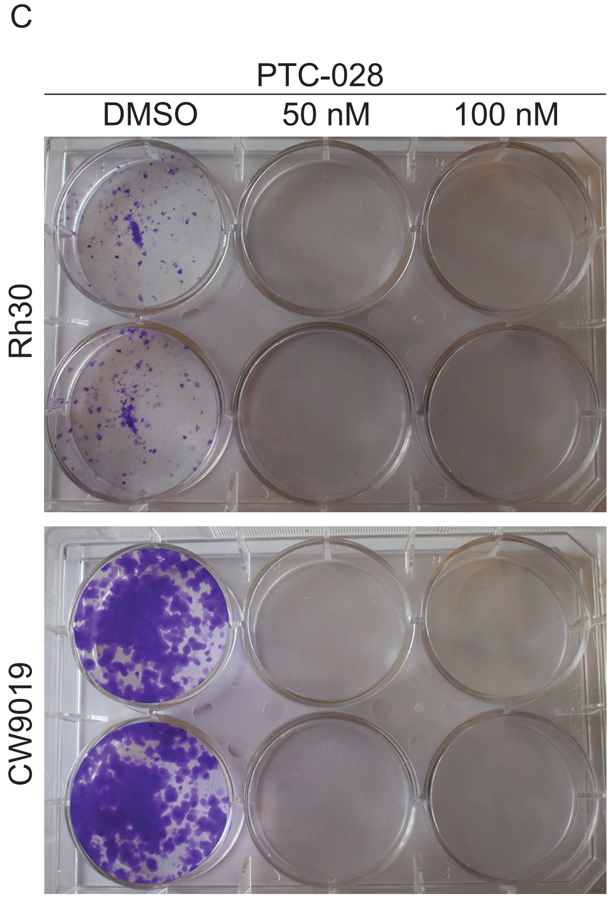

### Supplementary Figure S3

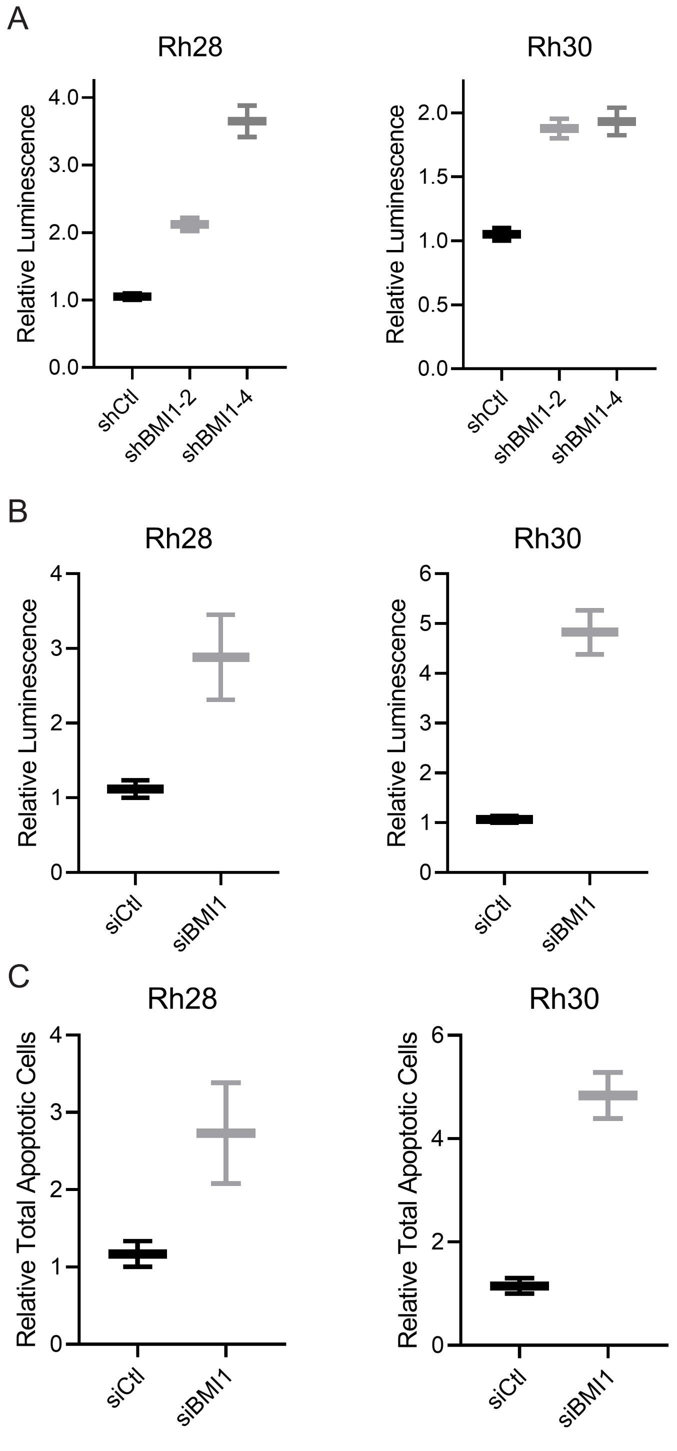

**Supplementary Figure S4**

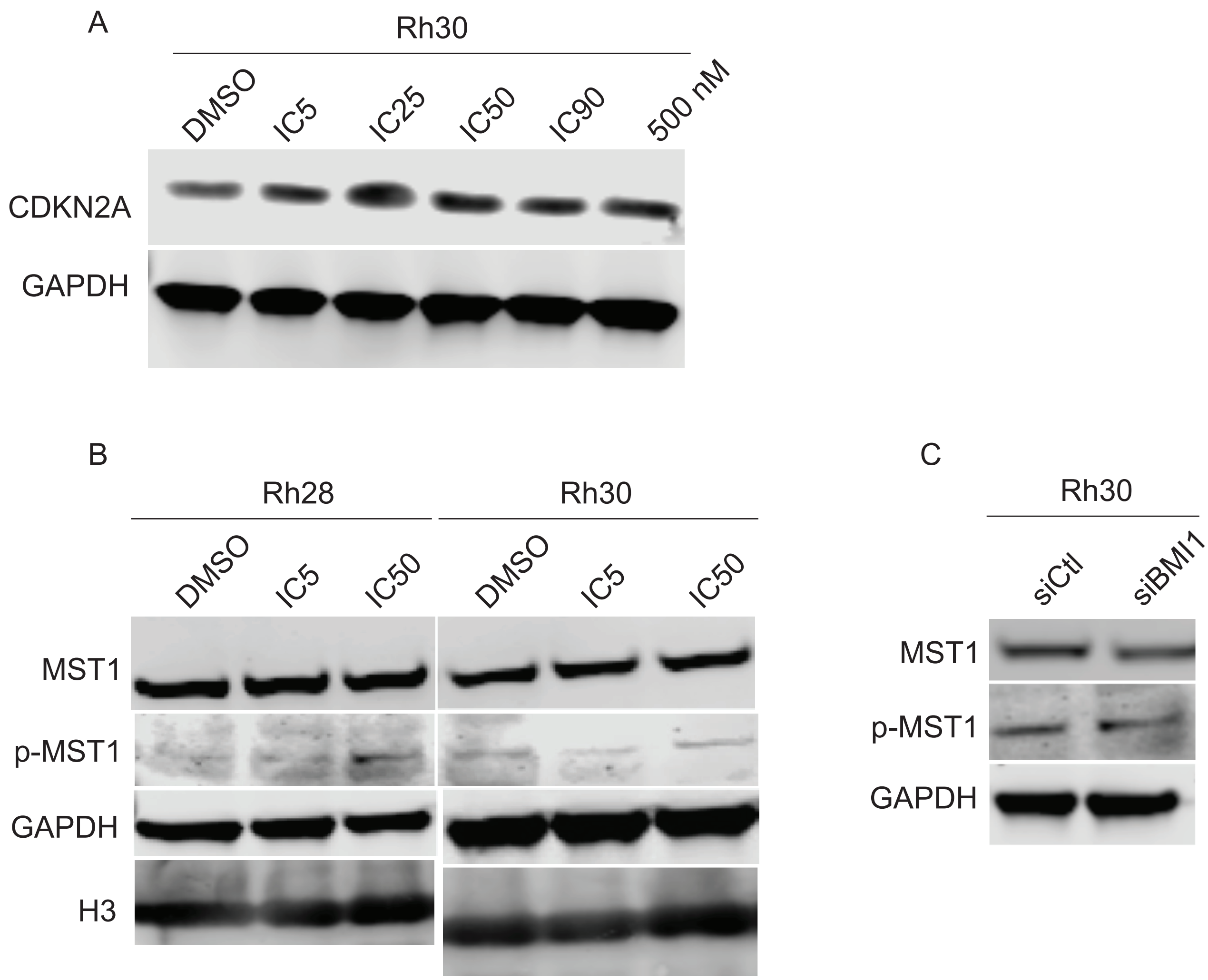
