## Supplementary Figure Legends for "Epigenetic regulator BMI1 promotes fusion-positive rhabdomyosarcoma proliferation and constitutes a novel therapeutic target"

**Supplementary Figure S1, related to Figure 1. *BMI1* is highly expressed in rhabdomyosarcoma**

(A-B) *BMI1* mRNA expression from Gibault (A) and Henderson Sarcoma (B). Human exome array data from the Oncomine database.^36^ Y-axis is Log2 median-centered intensity. (C) Data from the OncoGenomics database.^40^ *BMI1* expression on y-axis (Log2) from RNA-seq from FP-RMS and FN-RMS PDX models.

**Supplementary Figure S2, related to Figure 3. Pharmacologic inhibition of BMI1 decreases cell proliferation *in vitro***

(A) Rh30 cells were treated with a range of PTC-209 and PTC-028 doses and harvested after 72 hr. Western blots were performed and BMI1 levels were analyzed. Ku80 serves as a loading control. (B) Brightfield microscopy images (scale bar = 100 μm) of Rh30 cells treated with DMSO, 50 nM PTC-028, or 100 nM PTC-028 after 72 hr. (C) Images of crystal violet stained Rh30 or CW9019 cells treated with either DMSO or PTC-028 after 10 days of colony formation.

**Supplementary Figure S3, related to Figure 4. Targeting BMI1 decreases cell cycle progression and increases apoptosis in FP-RMS**

(A) Rh28 and Rh30 cells were transduced with two separate BMI1 (2 and 4) shRNAs compared to a nontargeting shRNA shCtl. Relative Caspase-Glo data measured in relative luminescence (y-axis) compared to shCtl. Western blotting as depicted in Figure 2A-B. (B) Rh28 and Rh30 cells were transfected transiently with an siRNA pool against BMI1. After 72 hr, the Caspase-Glo assay was performed and measured in relative luminescence (y-axis). RT-PCR as depicted in Figure 2C-D. (C) Annexin-V/PI staining was performed on siRNA transfected cells after 72 hr and total apoptotic cells were calculated. Standard deviation bars depicted. Experiments were performed in duplicate.

**Supplementary Figure S4, related to Figure 6. BMI1 negatively influences Hippo signaling**

(A). Western blot of CDKN2A levels across a range of PTC-028 doses in Rh30 cells. Cells were harvested after 72 hr. GAPDH as a loading control. (B). Western blot of MST1 and p-MST1 levels across a range of PTC-028 doses in Rh28 and Rh30 cells. GAPDH and H3 as loading controls. (C) Western blot of siRNA transfected cells of MST1 and p-MST1 levels in Rh30 cells. Western blotting of BMI1 as depicted in Figure 6C. GAPDH as a loading control.

**Supplementary Table 1**

List of antibodies, sources and dilutions used in all Western blot assays.

**Supplementary Immunoblots.** Original immunoblots are provided.
