## Supplementary Table 1 for "Epigenetic regulator BMI1 promotes fusion-positive rhabdomyosarcoma proliferation and constitutes a novel therapeutic target"

| **Antibody** | **Company** | **Dilution** |
| --- | --- | --- |
| BMI1 | Cell Signaling Technology | 1:1000 |
| Ku80 | Cell Signaling Technology | 1:5000 |
| Cleaved PARP | Cell Signaling Technology | 1:1000 |
| β-Actin | Cell Signaling Technology | 1:1000 |
| GAPDH | Cell Signaling Technology | 1:5000 |
| MST1 | Cell Signaling Technology | 1:1000 |
| p-MST1/2 (Thr183)/(Thr180) | Cell Signaling Technology | 1:500 |
| LATS1 | Cell Signaling Technology | 1:1000 |
| p-LATS1/2 (Thr1079)/(Thr1041) | Cell Signaling Technology | 1:500 |
| YAP/TAZ | Cell Signaling Technology | 1:500 |
| FOXO1 | Cell Signaling Technology | 1:1000 |
| CDKN2A (p16-INK4A) | Proteintech | 1:1000 |
| Histone H3 | Abcam | 1:1000 |

**Supplementary Table 1**
